# Exploring a large cancer cell line RNA-sequencing dataset with k-mers

**DOI:** 10.1101/2024.02.27.581927

**Authors:** Chloé Bessière, Haoliang Xue, Benoit Guibert, Anthony Boureux, Florence Rufflé, Julien Viot, Rayan Chikhi, Mikaël Salson, Camille Marchet, Thérèse Commes, Daniel Gautheret

## Abstract

Analyzing the immense diversity of RNA isoforms in large RNA-seq repositories requires laborious data processing using specialized tools. Indexing techniques based on k-mers have previously been effective at searching for RNA sequences across thousands of RNA-seq libraries but falling short of enabling direct RNA quantification. We show here that RNAs queried in the form of k-mer sets can be quantified in seconds, with a precision akin to that of conventional RNA quantification methods. We showcase several applications by exploring an index of the Cancer Cell Line Encyclopedia (CCLE) collection consisting of 1019 RNA-seq samples. Non-reference RNA sequences such as RNAs harboring driver mutations and fusions, splicing isoforms or RNAs derived from repetitive elements, can be retrieved with high accuracy. Moreover, we show that k-mer indexing offers a powerful means to reveal variant RNAs induced by specific gene alterations, for instance in splicing factors. A web server allows public queries in CCLE and other indexes: https://transipedia.fr. Code is provided to allow users to set up their own server from any RNA-seq dataset.

## 1 Introduction

RNA expression analysis now plays an essential part in molecular biology and medicine. Public RNA-sequencing (RNA-seq) repositories have grown in size to millions of samples, each several Gbytes in size. The Sequence Read Archive (SRA) alone contains 1.8 million public human RNA-sequencing experiments as of January 2024. Due to high costs of RNA-seq data download and reanalysis, exploration of RNA-seq repositories is typically confined to precomputed gene expression tables [1, 2]. As it is restricted to annotated genes or transcripts, this approach overlooks a large part of transcriptional diversity, which includes mutated, abnormally spliced, intergenic, intronic, repetitive, or fusion RNAs [3]. Projects such as Recount offer a way to query independent exons or splice junctions in very large (SRA-scale) datasets [4], however this still relies on sequence alignments and does not allow to quantify an arbitrary RNA directly. Considering the huge diversity of RNA forms, searching RNA-seq repositories using current tools is like looking under the proverbial lamppost. New methods are required to explore the hidden diversity within RNA-seq data.

Reference-free queries in large sequence sets are possible thanks to several k-mer based data structures that can index large sequence datasets in a fraction of the disk space used for raw sequences (see [5] for review). However, most k-mer data structures are limited to qualitative queries (presence or absence of a given sequence), which is not satisfying for RNA expression analysis. Three recent tools enable quantitative queries in large sequence sets. Needle [6] implements multiple interleaved Bloom filters and sketches of minimisers, which enable storing counts in a semi-quantitative way. Metagraph [7] uses an optimized De Bruijn Graph structure, enabling to store either presence-absence or count information. While Metagraph proposes ready-made indexes for diverse collections of genomes and metagenomes, the public server does not return count information and is limited to one query sequence at a time. Our indexing tool Reindeer [8] is optimized for processing several thousands of samples and associate k-mers to approximate but accurate counts in each sample.

Here we use an improved version of Reindeer deployed on a web server to demonstrate the capacity of reference-free RNA-seq indexes to detect and quantify arbitrary RNA variations of biological significance in cancer RNA-seq data. First we re-evaluate the computational time and memory footprint of Reindeer in this practical setting. We then show that transcript quantification with Reindeer can achieve a high accuracy by masking non-specific sequences in queries. Building upon this, we introduce the first public reference-free index of the CCLE RNA-seq database. The rich biological data in CCLE (1019 cell lines from 40 tumor types) allows us to illustrate Reindeer’s ability to accurately detect and quantify a large diversity of non-reference RNA sequences, including RNA mutations, fusions, transposable elements and splice variants. The reference-free CCLE RNA atlas is available for online queries along with other datasets at https://transipedia.fr.

## **2** Results

### Indexes for arbitrary RNA sequence query and quantification

Our objective is to provide a computational framework enabling quantification of arbitrary RNA sequences in large RNA-seq datasets. This framework must satisfy several criteria: (i) the capability to index any RNA-seq dataset while preserving all information at single-base resolution, and (ii) the ability to query the index in realtime for quantifying the occurrence of input sequences in each sample within the index. Indexes should be available for query either through a web interface or on a local computer. We describe below the realization of such a framework using Reindeer.

### Building and querying indexes

The implementation of a Reindeer index server is presented in Fig. 1A. Indexes were created with a k-mer size of 31, using the on-disk option that allows queries to be performed while only storing the primary k-mer hash in memory. Currently available online indexes cover 151 billion reads in 1851 samples. Indexes have relatively small memory footprints and file sizes 15 to 40 times smaller than the original compressed fastq files (Table 1). For instance, hardware requirements for querying the 1019-sample CCLE index are only 22.3Gb RAM and 236Gb disk. A socket mode enables the index to reside in memory once loaded and allows for real-time queries. Users submit queries through the web interface and receive results in the form of count tables or graphics (Fig. 1B). Query times are fast enough to handle mutiple interactive queries for short sequences or full-length mRNAs (100,000 31-mers in 16s, 100 mRNAs in 12.6s) (Table 2).

**Figure 1:**
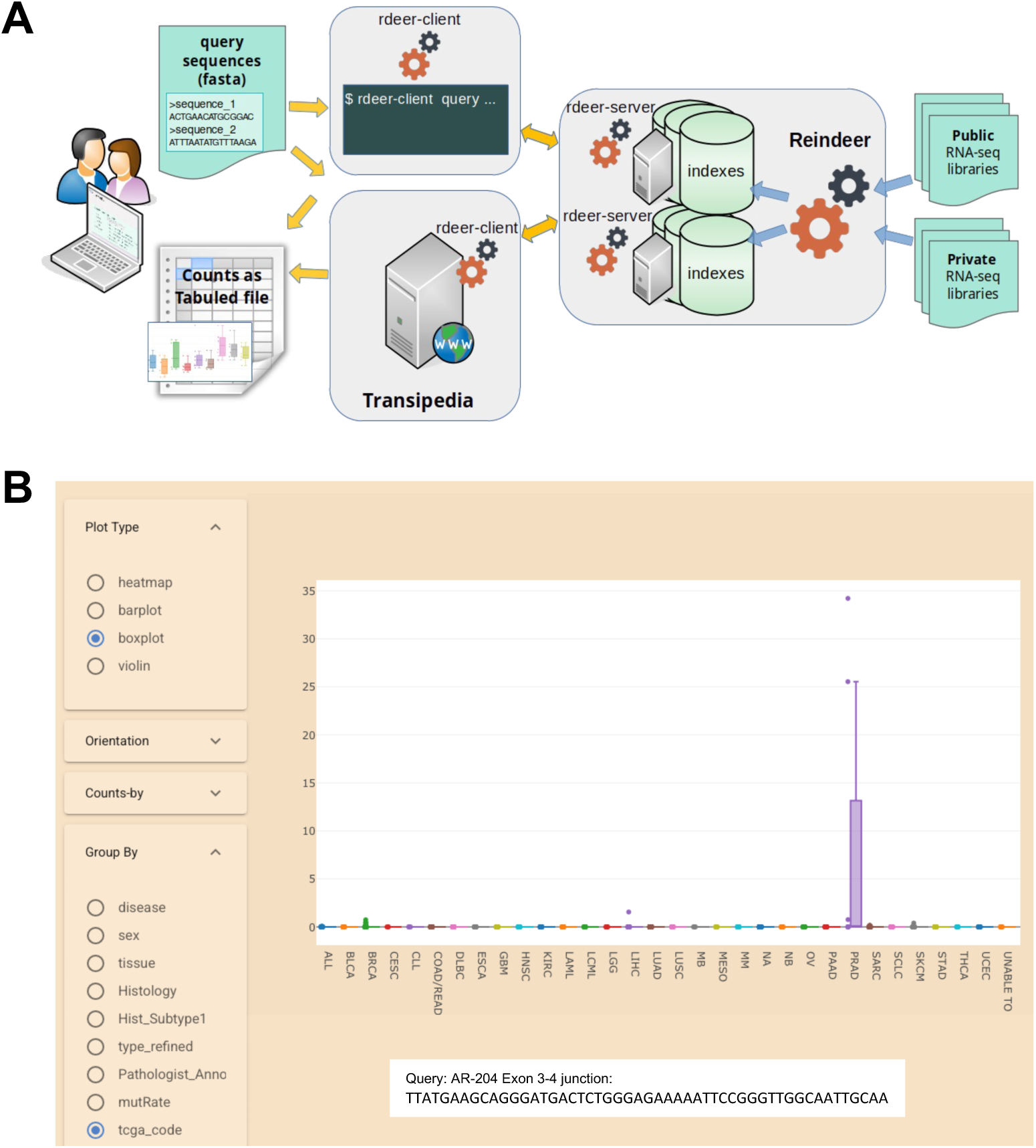
**A:** Reindeer index build and query workflow implemented on the Transipedia web server. Reindeer pre-built indexes can be easily queried by nonexperts using the Transipedia web interface. For massive query and/or pipelines, the index can be queried from the command line using rdeer-client (rdeer). The output is a tabulated count file. **B:** Graphical output of a query corresponding to a prostate cancer-specific sequence (AR-V7). Each dot corresponds to a cell line, with Reindeer counts on the Y axis. The query sequence was a 51-nt fragment spanning exon 3-4 junction, specific to androgen receptor variant AR-V7 (Gencode transcript AR-204). Examples of web queries are provided on the repository: https://github.com/Transipedia/Reindeer-use-cases

**Table 1:** Reindeer index properties for various datasets (on-disk indexes)

| Dataset | #Samples | Fastq.gz size (Gb) | Index size (Gb) | RAM (Gb) | Load time (h:m:s) |
| --- | --- | --- | --- | --- | --- |
| GSE62852-AML | 40 | 252 | 16 | 10.8 | 00:02:17 |
| GTEX (part) | 1119 | 6100 | 312 | 42.2 | 00:08:58 |
| SEQC/MAQC | 16 | 51 | 2.4 | 3.1 | 00:00:33 |
| CCLE | 1019 | 8900 | 236 | 22.3 | 00:05:57 |

**Table 2:** query times on the CCLE index (on-disk index)

| Query type | # Queries | Query time (sec) |
| --- | --- | --- |
| 31-mers | 1000 | 1.0 |
|  | 10000 | 2.0 |
|  | 100000 | 16.0 |
|  | 500000 | 89.0 |
|  | 1000000 | 179.0 |
| full-length mRNAs<br>(mean size: 1.9kb) | 1 | 0.6 |
|  | 100 | 14.5 |
|  | 1000 | 132.8 |

### Accuracy of RNA expression measure

In order to assess Reindeer’s capacity to accurately quantify RNA expression from RNA-seq samples, we compared it to standard quantification approaches. Reindeer queries can be made using full-length sequences (e.g., complete mRNAs) or fragments of size not smaller than *k* as input. Reindeer returns counts for all consecutive k-mers in the query (Fig S1, suppl. material). Counts can be interpreted in different ways depending on whether users expect raw counts or counts normalized by query sequence length. To determine the optimal counting scheme, we used the SEQC/MAPQC dataset in which the abundance of 1000 transcripts was evaluated in 16 samples both by qPCR and Illumina RNA-seq [9]. Means of k-mer counts best correlated with qPCR abundance and transcript-per-million (TPM) measured from RNA-seq reads by Kallisto [10] (Fig 2A,B, Fig S2), while sums of k-mer counts best correlated with raw RNA-seq counts (Fig 2C, Fig S2). Correlation coefficients (CC) with Kallisto counts were around 0.8, in line with previous reports [6]. We found that quantification accuracy could be substantially improved by masking query k-mers with multiple instances in the human genome (Methods). This procedure led to *>*0.9 correlations with both qPCR and RNA-seq derived abundances, reaching a Pearson CC of 0.95 with Kallisto raw counts (Fig 2D-F, Fig S2). This demonstrates that simple quantitative queries in a k-mer index can achieve accuracies approaching that of a state-of-the-art RNAseq quantification method. Note that while TPM-like counts are identical in absolute value across methods, raw counts require a linear correction due to the conversion of fragment to k-mers (discussed in suppl. material).

**Figure 2:**
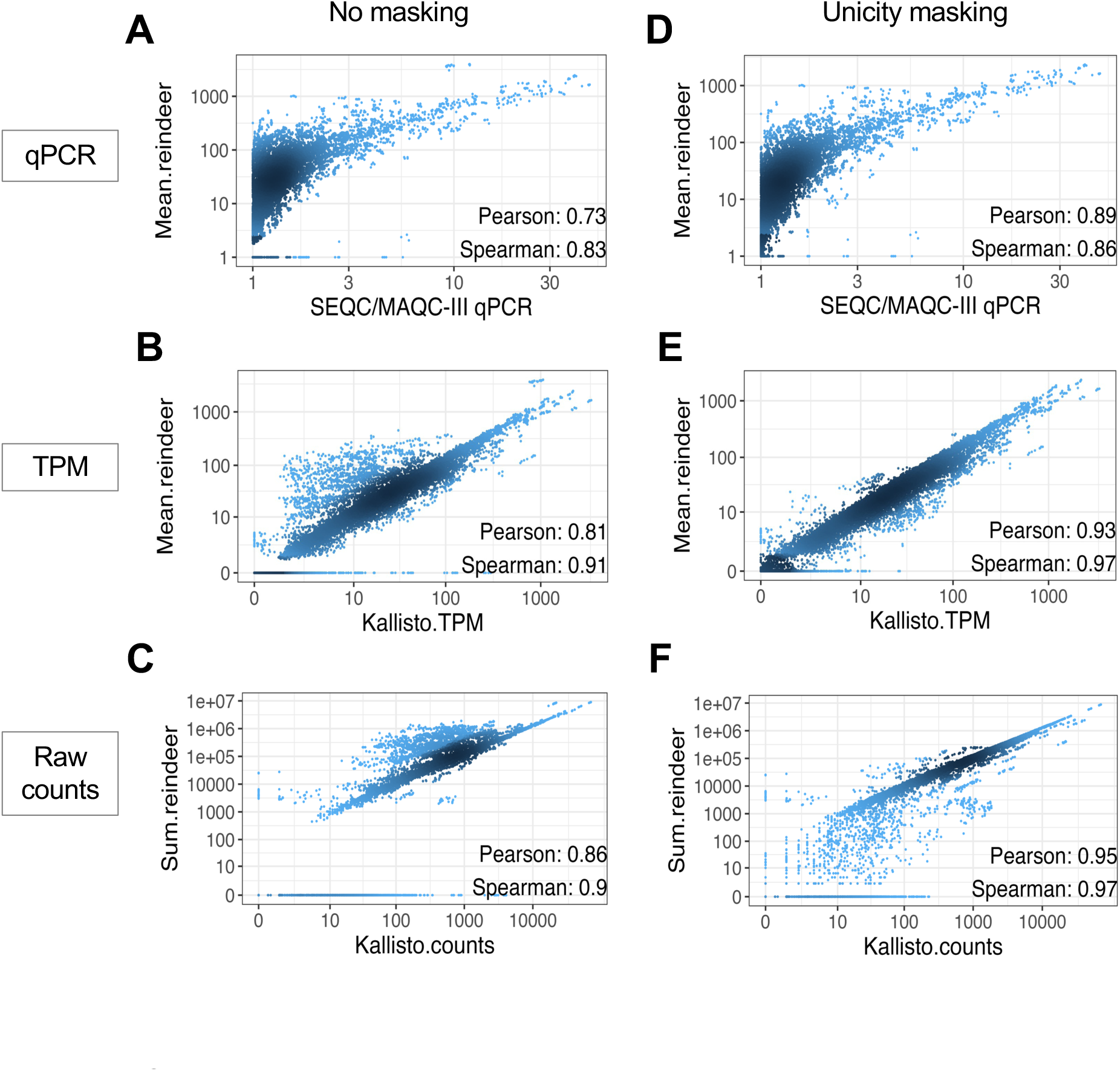
Correlations between Reindeer counts and established count methods. 1000 genes were quantified in 16 reference SEQC/MAQC-III samples. A,D: Reindeer (mean counts) vs. qPCR; B,E: Reindeer (mean counts) vs. Kallisto TPM; C,F: Reindeer (sum counts) vs Kallisto raw counts. Unicity masking: counts obtained after removal of non-unique k-mers.

### Finding mutations in RNA

Given an index of 1019 cancer cell lines enabling fast and accurate quantification of arbitrary RNAs, we set out to use this system to retrieve different types of RNA variations not commonly accessible in transcriptome databases. First we designed queries for mutations and indels. We selected mutations/indels in common cancer genes from the Depmap database [11] and designed 61-nucleotides sequences around each variation as explained in Methods (Fig. S3, Table S1). We refer to these sequences as ”probes”. With *k*=31, a 61-nucleotides (2*k* − 1) size ensures that any k-mer in the probe covers the variation.

To limit false positive calls, we applied a masking step that discarded parts of probes with multiple hits in the genome or harboring low complexity sequences (Methods). While this only eliminated 3.2% of probes (Table 3), it reduced false positive hits by 93.9% (Table S2). Several query modes were then tested whereby at least 1, 3, 5 or 10 k-mers in each probe had to be non-zero for the call to be made (*min hits*=1 to 10)(Table 3). Recall was satisfying in all cases (0.88 to 0.95) while precision ranged form 0.269 (*min hits*=1) to 0.89 (*min hits*=10). Thus, there is a significant benefit in requiring several k-mer hits around an event to make a call. Hereafter, *min hits* is set to 3 unless specified otherwise.

**Table 3:** Accuracy measures of Reindeer mutation and fusion calls.

|  |  | #total probes | #probes after selection<sup>1</sup> |  | *min_hits*<sup>3</sup>=1 | *min_hits*=3 | *min_hits*=5 | *min_hits*=10 |
| --- | --- | --- | --- | --- | --- | --- | --- | --- |
| Mutations | All Depmap mutations<sup>2</sup><br>(50 cancer genes) | 3685 | 3621 | <sup>4</sup> True + | 4346 | 4255 | 4205 | 4026 |
|  |  |  |  | False + | 11810 | 911 | 589 | **484** |
|  |  |  |  | False - | **255** | 346 | 396 | 575 |
|  |  |  |  | Precision | 0.269 | 0.824 | 0.877 | **0.893** |
|  |  |  |  | Recall | **0.945** | 0.925 | 0.914 | 0.875 |
|  | Cosmic Hotspot mutations | 960 | 951 | True + | 1665 | 1631 | 1611 | 1558 |
|  |  |  |  | False + | 6823 | 184 | 114 | **90** |
|  |  |  |  | False - | **51** | 85 | 105 | 158 |
|  |  |  |  | Precision | 0.196 | 0.899 | 0.934 | **0.945** |
|  |  |  |  | Recall | **0.970** | 0.950 | 0.939 | 0.908 |
| Fusions | All Depmap fusions<br>(junction at exon edges) | 8972 | 8860 | True + | 9410 | 9277 | 9201 | 9018 |
|  |  |  |  | False + | 25732 | 10048 | 6558 | **2378** |
|  |  |  |  | False - | **170** | 303 | 379 | 562 |
|  |  |  |  | Precision | 0.268 | 0.480 | 0.584 | **0.791** |
|  |  |  |  | Recall | **0.982** | 0.968 | 0.960 | 0.941 |
|  | Cosmic fusions | 60 | 59 | True + | 99 | 98 | 98 | 96 |
|  |  |  |  | False + | 22 | 3 | 3 | **2** |
|  |  |  |  | False - | **1** | 2 | 2 | 4 |
|  |  |  |  | Precision | 0.818 | 0.970 | 0.970 | **0.980** |
|  |  |  |  | Recall | **0.990** | 0.980 | 0.980 | 0.960 |

<sup>1</sup>: Min count cutoff and low complexity masking
<sup>2</sup>: Restricted to RNA-seq-derived mutations
<sup>3</sup>: *min\_hits*: minimum number of positive k-mers in query
<sup>4</sup>: All positive and negative counts are given for pairs {probe, sample}

Restricting Reindeer queries to established cancer-related (”hotspot”) mutations from Cosmic [12] substantially improved precision and recall (≥ 0.9, Table 3). We hypothesized that the remaining false positive (FP) calls may be true mutations filtered out by Depmap due to a more stringent count threshold. To assess this, we computed the variant allele frequencies (VAF) of mutations using counts obtained with wildtype and mutant probes. VAF computed by Reindeer was in general highly correlated to that inferred from conventional RNA-seq alignment (Fig. 3A) and FP calls had significantly lower VAF (Table S3, Fig. 3B), supporting these may be in part censored by Depmap. Further testing of 12 RNA-seq files (corresponding to 77 FP pairs) using a sensitive variant caller [13] or by direct parsing of the fastq files (linux *grep* command) confirmed 45 of the 77 (58%) of the putative FPs as likely true positives (Table S4). Finally, we analyzed samples with available DNA sequencing data: out of 44 FP calls in these samples, 31 (70%) turned out positives at the DNA level (Table S5). In summary, we estimate that more than half of the putative FP mutations at *min hits*=3 are actually true mutations.

**Figure 3:**
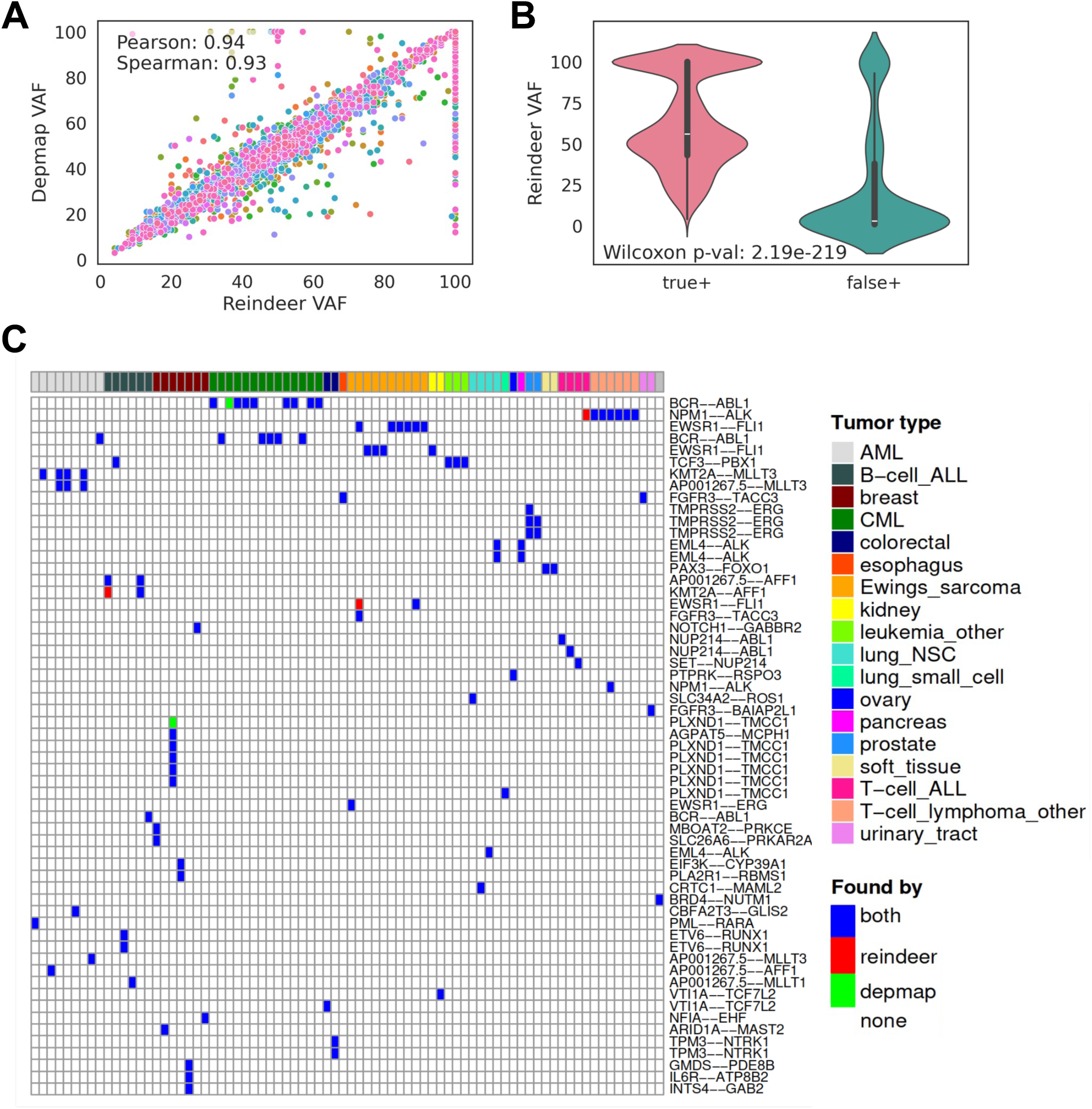
Comparison of Depmap and Reindeer calls for mutations and fusion transcripts. **A-B:** Reindeer variant allele frequencies (VAF) measured as the count ratio: mutated / (mutated + wild-type) * 100, for all Depmap mutations in cancer genes. **A:** Correlation of Depmap VAF (based on RNA-seq alignment) and Reindeer VAF. Each dot shows a mutation in one sample, colored according to gene (50 genes). **B:** Comparison of Reindeer VAF for true positive (n = 4255) and false positive (n = 911) calls in Depmap. **C:** Detection of DepMap Cosmic fusion events in CCLE cancer cell lines. Cosmic fusions were retrieved using a 51 nt probe centered on the fusion junction. Top: cell lines are colored by tumor type. Blue: events from DepMap found by Reindeer (True Positive); red: events found in an extra sample with Reindeer compared to DepMap; green: events not found with Reindeer. Lines with identical fusion names correspond to different exon-exon junctions of the same genes.

### Finding fusion transcripts

We next tested Reindeer’s capacity to retrieve gene fusion events. DepMap provides genomic coordinates of fusion junctions identified after alignment of RNA-seq reads by STAR-fusion [14]. We selected fusion events with a breakpoint at exon edges, which are considered more reliable [15], and designed 51-mer sequences centered on the fusion junction (Methods, Fig. S3B & Table S6). Probes shorter than (2*k* − 1) are desirable when querying fusion and splice junctions, since k-mers overlapping the junction at their tip might accidentally match other partner exons. Masking of k-mers present in the reference genome or transcriptome and of low complexity k-mers (see Methods) yield a total of 8860 fusion probes (Table S2).

Fusion events were quantified requesting at least 1, 3, 5 or 10 non-zero count k-mers, as done for mutations (Table 3). Recall was high in all cases (0.94 to 0.98), but precision was relatively low (0.27 to 0.79) due to a high number of FPs. Restricting evaluation to Cosmic fusions (100 fusion events) largely reduced the FP rate, improving both precision and recall to above 0.97 for *min hits >*= 3 (Table 3, Fig. 3C). This suggests the initial query list from Depmap contained fusions yielding multiple erroneous hits. The only two missed fusion events had SNPs in close proximity (7 and 4 nucleotides) to the junction, such that the minimum number of matching k-mers was not reached (Fig S4). Of the three apparent false positives remaining (Figure 3C, red), two were annotated in the LigeA fusion database [16] in the correct cell line, supporting their reality. Finally, fusion transcript expression quantified by Reindeer was highly correlated to that given by Depmap (Pearson CC=0.92, Fig S5).

### Finding expressed transposable elements

Transposable elements in the human genome are mostly silent but can be re-expressed in tumor cells upon lifting of epigenetic repression. Measuring their expression is complex because exact repeats impedes the attribution of RNA-seq reads to specific loci. We compared the quantification of human endogenous retroviruses (ERV, a major class of transposable elements) by Reindeer and by two software relying on different mapping strategies. Telescope [17] estimates transposable element expression at locus-level through genome mapping, allowing for up to 100 mapping positions and reassigning ambiguous reads to specific loci using an expectation maximization algorithm. While Reindeer does not use expectation maximization, locus-level ERV quantification after masking of non-unique sequences was reasonably similar to that of Telescope (Pearson CC:0.88, Fig. 4A, Fig. S6), while requiring only a fraction of the time (4-5 hours by sample with Telescope *vs.* seconds for Reindeer). REdiscoverTE [18] estimates transposable elements expression at the family level based on Salmon [19], a fast quantifier using pseudo-mapping. REdiscoverTE and Reindeer ERV quantifications were highly correlated, both for raw counts and for CPM normalized counts (Pearson CC=0.99 and 0.96 respectively) (Fig. 4B, Fig S7). Although a significant fraction of k-mers in ERV elements were masked as non-unique (Methods), all tested elements had sufficient specific k-mers to remain quantifiable, even at the locus level. Finally, as observed for mRNA quantification, Reindeer’s raw counts required a linear correction to match raw counts from the specialized tools.

**Figure 4:**
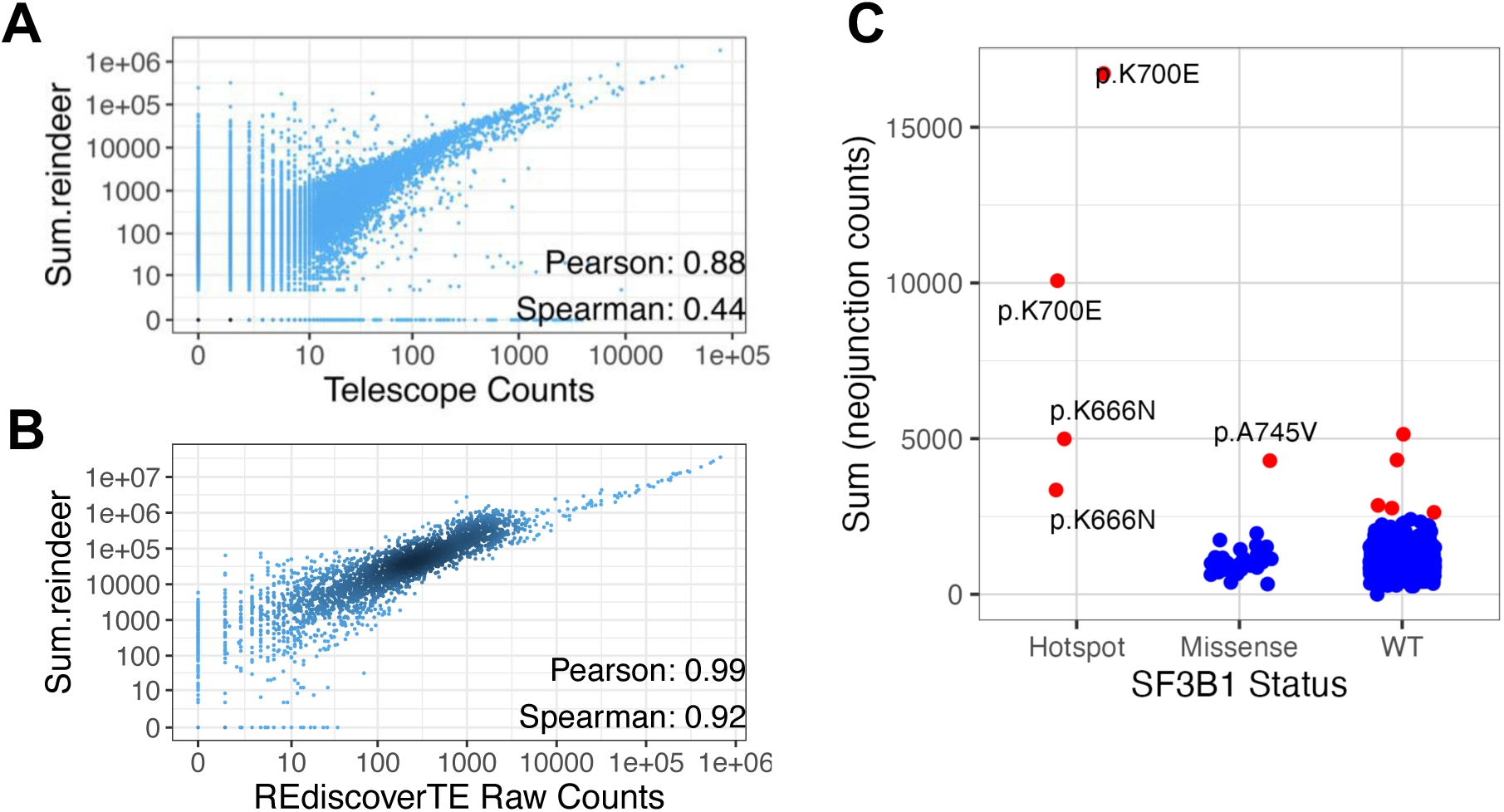
Quantification of transposable elements and novel splice junctions. Quantification of transposable elements and novel splice junctions. **A**: Correlation of quantification of 1000 ERVs by Reindeer and Telescope, in 57 colon cell lines from CCLE. **B**: Correlation of quantification of 50 ERV families by Reindeer and RediscoverTE, in 57 colon cell lines from CCLE. **C:** Quantification of SF3B1-induced neojunctions in cell lines. Each dot represents the sum of counts of 849 SF3B1-induced neojunctions in one CCLE cell line. Cell lines harboring hotspot (oncogenic) SF3B1 mutations, other missense SF3B1 mutations and wild-type SF3B1 are distinguished. Cell lines with outlier neojunction expression (>2 SD above mean) are shown in red.

### Finding aberrant splicing junctions

Aberrant splice junctions caused by mutations in RNA processing genes are generally absent from reference transcriptomes. Their detection usually requires downloading and reanalyzing RNA-seq files. Using Reindeer, one may directly interrogate an RNA-seq index for such unreferenced variants. We illustrate this with splicing alterations in uveal melanoma. Mutation of the SF3B1 splice factor in uveal melanoma induces aberrant splicing of hundreds of genes [20]. We retrieved aberrant splice junctions observed in SF3B1-mutated patients and created 51 nucleotides probes for 849 so called neojunctions (see Methods). These sequences were then quantified in CCLE (Fig. 4.C). All cell lines harboring known oncogenic SF3B1 mutations presented significantly elevated neojunction expression, consistent with genome-wide SF3B1-induced alterations. Another SF3B1 mutation with elevated neojunctions was A745V in NCIH358 LUNG, suggesting this mutation may also disrupt splicing, although this is not documented in the current litterature. Moreover, five cell lines with no SF3B1 mutation behaved like SF3B1 hotspot mutants, suggesting alterations in the same splicing pathway in these cells. This included two lung and two endometrial tumors, which are tumors where SF3B1 and related SUGP1 mutations are documented [21, 22]. Interestingly, two of these cell lines had impairing mutations in SUGP1 (Table S7). This illustrates how a Reindeer index can be utilized to evaluate a complex transcriptome signature composed of aberrant transcripts and identify cells altered in similar pathways.

## **3** Discussion

We describe here the first practical implementation of a web server for reference-free, quantitative queries in an RNA-seq dataset of over 1000 samples. The service runs on a standard computer using less than 25Gb memory and 250Gb SSD storage. It was tested with a variety of input queries including full-length mRNAs and RNA elements that are not usually represented in curated RNA-seq databases, such as transposable elements, fusions, neo-splice junctions and mutated RNAs. When using the system to retrieve known mutation and fusions events, precision and recall were above 0.9 for oncogenic events. Furthermore, a large fraction of inferred false positives were shown to be likely true events filtered out in the reference database.

Reindeer count accuracy was high in spite of the conversion of read counts into aggregated k-mer counts at indexing, and their subsequent conversion to query-level counts when processing query results. Count correlation with state-of-the-art quantification methods were always above 0.8 (Pearson CC) for full-length mRNAs, fusion transcripts and transposable elements, and above 0.9 after masking non-specific k-mers from queries.

A lesson learnt during this study was the importance of ”query engineering”. The best accuracy was reached through proper query design, masking and post-processing. Query design involves selecting the right ”probes” to ensure returned hits do not include unspecific sequences. With our default k-mer size of 31 nucleotides, optimal probes were 61nt fragments around mutations or 51nt fragments around splice or fusion junctions. Query masking involved removal of non-specific (non-unique and low complexity) k-mers from queries. This provided important gains in count accuracies for all types of queries. Furthermore, this considerably reduced the number of false positives when querying local events such as mutations. The query design and masking methods introduced herein could serve as guidelines to users of k-mer based indexes in general.

Post-processing of query results first involves deciding how many k-mers in a query must be matched to accept a hit. This step is only important for local event detection (mutations, fusions, splice junctions), in order to accomodate possible SNPs around events. We identified the optimal setting whereby SNPs were almost never a problem while retaining a high specificity. The second post-processing step is the conversion of Reinder k-mer counts into TPM-like or raw-count-like values. Averaging k-mer counts provided count estimates that were remarkably similar to TPM, while summed counts were highly correlated to raw counts, albeit with a conversion factor.

Some limitations of the current Reindeer framework must be acknowledged. (i) Reindeer index building is a separate action that is computer intensive and best performed by bioinformaticians. (ii) Real-time queries are available to web users thanks to preloaded indexes. Tools for pre-loading indexes are provided in Methods. However, local instances will have to load indexes into memory first, which may take several minutes before queries are processed. (iii) Query design may require running an independent tool such as the *Kmerator Suite* [23] prior to submitting queries. This may be further integrated into the server after enough user experience is gathered.

## **4** Conclusion

Reference-free indexes provide a direct access to unprocessed RNA-seq data, enabling biologists to ask questions that would otherwise require resource-intensive pipelines. Beyond obvious applications such as verifying the tissue or tumor specificity of novel biomarkers, Reindeer’s quantitative indexes allow to carry out sophisticated experiments by simultaneously querying oncogenic alleles, RNA isoforms, repeats, etc. and process the resulting count table to uncover novel functional interactions. We hope to expand the Transipedia server to include an increasing number of public datasets to facilitate this type of experiment.

## **5** Methods

### Updates to Reindeer

Since its initial publication [8], Reindeer has been enhanced with a socket mode to facilitate remote server queries. This improvement enables the efficient management of indexes from various collections and ensures rapid query responses. Reindeer utilizes an effective k-mer hashing structure to map k-mers to their respective counts in each sample, alongside a matrix that represents the abundance of indexed objects across samples. Through extensive testing, we observed that the primary bottleneck in many use cases was the loading of the index into RAM, while the actual querying process is quickly expedited thanks to the hashing structure. As a result, Reindeer’s default algorithm was transitioned from relying predominantly on in-RAM queries to disk-based queries. This shift involves the ability to serialize the count matrix of Reindeer onto the disk in a compressed format. Consequently, we updated Reindeer to only load the hashmap into RAM in the initial phase, markedly reducing the total time required for conducting intensive queries, especially when running on SSD.

### Building and using Reindeer indexes

Building codes for the web and local server environment are described in https://github.com/Transipedia/publication-ccle. RNA-seq data source is described in Table S8. CCLE RNA-seq raw fastq files were retrieved from Gene Expression Ominibus dataset GSE36139. Fastq files were first checked for sequence quality using FastQC (version 0.11.9), MultiQC (version 1.9) and KmerExplor [23] for contaminations and library information. Cutadapt (version 1.18) was used for low-quality trimming (-q 10,10), excluding sequences shorter than 31 nt after trimming (-m 31). Adapter sequence removal was deemed unnecessary in the studied datasets. Fastq files were then processed by bcalm v2.3.0 (https://github.com/GATB/bcalm). For the CCLE dataset, k-mers with counts *<* 4 were excluded (option -abundance-min 4). Bcalm files were then used as input to Reindeer v1.02 (https://github.com/kamimrcht/REINDEER). Indexes for the web server were built using the on-disk option. For querying, indexes were copied to a SSD drive (applies to the web server too). All query times were obtained using the rdeer-client software running on a local index and include count aggregations for multi-probe queries.

Gene expression quantification benchmark.

The SEQC/MAQC-III dataset [9] provides both RNA-seq and qRT-PCR values for 1000 genes across 16 reference samples. We used the 16 Illumina files and the preprocessed Taqman-raw.txt file from Chisanga et al. [24], retrieved from https://github.com/ShiLab-Bioinformatics/GeneAnnotation. RNA-seq data was processed as above. Gene expression was quantifed with Kallisto (version 0.46.1) using the v108 Ensembl transcriptome (cdna+ncrna), followed by tximport [25] for computing gene-level raw counts and TPM values. The Reindeer index was generated from trimmed fastq files using cutadapt -q 10,10 -m 31. Reindeer gene expression estimates were obtained using Ensembl v108 canonical transcripts as input, subject to the following processing steps.

### Query preparation

Queries were pre-processed to remove non-specific and low complexity k-mers. When ”masking” is specified, non-specific parts of query sequences were deleted using the Kmerator software (https://github.com/Transipedia/kmerator) [23]. Kmerator takes as input a genome index and a fasta file of query sequences or a list of gene names. By default any k-mer in the query that is present more than once in the genome is deleted. The optional parameter --max-on-transcriptome X requires that k-mers be present at most *X* times in the transcriptome annotation file (Ensembl v108 was used).

Low complexity masking discards k-mers meeting any of the following conditions:

- containing a *>*= 6−nt homopolymer, or
- 3-mer complexity defined as ((number of distinct 3-mers) / (total number of 3-mers)) below 0.55. This cutoff was determined from the analysis of complexity distribution in four independent datasets (Fig. S8).

### Processing of query results

Reindeer queries return a series of triplets *b_i_* − *e_i_* : *q_i_*, each corresponding to a monotig (Fig. S1 and suppl. material) matched by the query sequence. A ∗ symbol for *q_i_* means that the monotig does not have enough k-mers (with non-zero counts) for reporting a reliable result. This minimum k-mer presence criteria is provided as a percentage in the −*P* parameter. The default −*P* value (40%) was used unless otherwise specified. Query abundance in Fig. 2A-C was computed as the mean, median, maximum and sum values of monotig counts. Mean, median and sum were weighted by the number of k-mers in each monotig. The maximum value was calculated in the trivial way as it is not affected by k-mer multiplicities. For masked queries (Fig. 2D-G), substrings were queried separately by Reindeer and the resulting counts were merged per original query (this option is available on the web server, however it is only possible with *mean* abundance counting).

### RNA mutations

We selected 50 highly mutated cancer genes from CCLE [26], TCGA [27] and hematological malignancies [28] (Table S9). RNA-seq derived mutations within these genes were retrieved from the DepMap Public 22Q2 MAF mutation file (file CCLE mutations.tsv, field RNAseq AC) and converted into VCF format. This represented 3685 mutations, herein referred to as ”all Depmap”. A subset of 961 Hostpot mutations (*i.e.* known cancer drivers) was further selected based on field field CosmicHotSpot in the mutation file. The probe selection process for mutations is described in Figure S3A. For each mutation, a 61 nt-long probe centered on the mutation and its wildtype 61 nt-long counterpart were produced with *vcf2seq*. Probes were masked using Kmerator with the --chimera option that deletes any probe k-mer present in the reference genome or transcriptome for the mutation probes and with the --max-on-transcriptome 100 option that only deletes non-unique k-mers on the genome and *>*100 occurences on the transcriptome for the wild-type probes.

Probes were also masked for low complexity elements as described above. At the end of the masking process, about 3% and 1% of probes were deleted from the ”all Depmap” and hostpot probe sets, respectively (Table 3).

### Fusions

Fusions were retrieved from the DepMap Public 22Q2 fusion table (field CCLE fusions.csv). We set the minimum read count supporting a fusion to 4 (same as used in the Reindeer index) which retained 14946 fusions. A bed file was generated for the left and right sides of junctions and 51 nt-long probes centered on the junction were produced using bedtools getfasta[29]. We then selected fusions with junctions at exon edges by intersecting fusion coordinates with Gencode V42 exon coordinates (8972 fusions). K-mers were masked using kmerator --chimera and low complexity filter as above. The complete procedure is shown in Figure S3B. A total of 8860 fusion queries were eventually retained. A subset of 60 known oncogenic fusions was selected based on the ”Cosmic” label in column annots of the DepMap table. Selected fusions were further verified on the Ligea dataportal (http://hpc-bioinformatics.cineca.it/fusion/) which provides fusions predicted in CCLE RNA-seq data by four detection algorithms and enables retrieval of the corresponding read sequences.

### Transposable element expression

Transposable elements quantification was performed in 56 CCLE samples from colon cell lines. For comparison with Telescope [17] (V.1.0.3), we selected 1000 ERV loci (4034 sequences) from the authors’ supplemental data. We then generated query sequences based on genomic coordinates (Hg38), and masked non-unique sequences using kmerator with option --max-on-transcriptome 100. Unicity masking deleted 17% of k-mers in ERV probes in average. Nonetheless, every locus retained at least one probe with enough specific k-mer to be measurable. Telescope runtime was estimated based on a run with 16 threads and 48Gb RAM. For tests against REdiscoverTE [18], we retrieved genomic locations for 58 ERV families from the REdiscoverTE repository https://github.com/ucsffrancislab/REdiscoverTE/. This represented 40,734 loci, which were converted to sequences using bedtools, and masked for non-unique k-mers as above, resulting in 305,331 probes. Counts were aggregated at the family level.

### Neo-splicing events

The coordinates of 1258 abnormal splice junctions associated to SF3B1 mutations were retrieved from Table 2 of [20], converted to bed format and lifted to Hg38 using overlift (UCSC tools). As in the fusion procedure, we generated a 51-nt long sequence centered on the splicing junction and masked any genome or transcriptome k-mer (kmerator --max-on-transcriptome 0) and low complexity k-mers, retaining 849 probes. Probes were quantified in CCLE using the mean method.

### Declarations

- Funding: Agence Nationale de la Recherche grants ANR-18-CE45-0020, ANR-22-CE45-0007, ANR-19-CE45-0008, PIA/ANR16-CONV-0005, ANR-19-P3IA-0001.

This project has received funding from the European Union’s Horizon 2020 research and innovation programme under the Marie Sk-lodowska-Curie grants agreements No. 872539 and 956229.

- Conflict of interest/Competing interests: no competing interest to declare
- Ethics approval: not applicable
- Consent to participate: not applicable
- Consent for publication: all authors agreed with publication
- Availability of data and materials: see Methods.
- Code availability: https://github.com/Transipedia/publication-ccle
- Authors’ contributions: TC and DG supervised the study and wrote the manuscript. CB, TC, DG, RC, M and CM contributed to the study design. CB and HX performed analyses and co-wrote the manuscript. BG and AB developed interfacing utilities for Reindeer. FR and JV performed some of the analyses.

## Supplementary Information

### Definitions

#### k-mer

k-mers are short sequences in a fixed length k. In this article, we limit our scope on 31-mers formed by A, C, G and T, targeting for RNA-seq study.

k-mer count in RNA-seq sequence library. Given a k-mer and an RNA-seq library, the k-mer’s count is defined as its number of occurrence in the set of sequencing reads.

Sample count vector of a k-mer. Given a k-mer and *M* ordered RNA-seq libraries, the k-mer’s sample count vector is defined as an integer vector in an *M* -dimensional space, of which the *i^th^* component represents the k-mer’s count in the *i^th^* sequencing library (*i* = 1, 2, *…, M*).

Arbitrary sequence. An arbitrary sequence is formed by a finite, variable number of elements selected from a finite set, arranged in an arbitrary order. In the scope of this article, the sequence is considered as RNA sequences, formed by nucleotides A, C, G and T.

In confusion-free context, we simplify the arbitrary sequence as sequence to be consise.

Sequence query. Given an arbitrary sequence and a series of sequencing libraries, the task of arbitrary sequence query aims to either (i) qualitatively detect its presence/absence, or (ii) to quantitatively estimate its expression level, in each of the sequencing library. In this article, we address to the quantification aim, and we suppose the length of the arbitrary sequence is no less than 31 nucleotides (for 31-mers being used).

Component k-mers of a sequence. A sequence of length *L* can be destructed into (*L* − *k* + 1) consecutive k-mers, shifting one nucleotide by step. The set of (*L* − *k* + 1) k-mers form the sequence’s component k-mers.

Monotig Given a sequence, a monotig is one of its substring that has a uniform sample count vector across component k-mers.

Please note that: (i) a monotig is also an arbitrary sequence *per se*; (ii) unlike k-mers, different monotigs of a same arbitrary sequence do not overlap each other.

Monitigs are the elementary units for Reindeer query. Given a query sequence *Q*, one its monotig *m* is determined by a pair of positions *b_Q_*(*m*) and *e_Q_*(*m*), respectively meaning the starting and ending k-mers in the sequence *Q*. In the context where the query sequence *Q* is defined, we simplify the writing from *b_Q_*(*m*) to *b_i_* and *e_Q_*(*m*) to *e_i_*, where *i* is the monotig’s ordinal number along *Q*, i.e., determined as between the *b^th^* and *e^th^* k-mers along *Q*.

A monotig of a given query sequence can or cannot be found in one specific library *S*. A successful query is gained when the monotig has sufficient proportion of component k-mer counts in the *S* to support the hit. The minimum proportion is set by user with -P parameter of the software.

The count of a given monotig *m* in a library *S* is reported by a triplet *b_i_*-*e_i_*:*q_i_*, in which *b_i_*-*e_i_* specifies the starting and ending k-mers along the query sequence for *m*, and *q_i_* indicates the query result of the monotig’s component k-mers in library *S*. In the case of successful query, *q_i_* is returned as an integer, while when the query is failed, it is originally returned as an “*” sign by the software, and we replace the “*” sign by 0 in this article.

When Reindeer queries a sequence composing *n* monotigs, it returns for each library *S* a list of triplets indicating monotigs’ query results, formed as *b*_1_ − *e*_1_ : *q*_1_, *b*_2_ − *e*_2_ : *q*_2_, . . ., *b_n_* − *e_n_* : *q_n_*, where each triplet *b_i_* − *e_i_* : *q_i_* reports the count of the *i^th^* monotig (*i* = 1, 2, *…, n*).

### Conversion from read to k-mer counts

There is a difference in scale between Reindeer sum counts and raw counts from the various quantification tools (Fig 3C, Fig. 4A,B). The average correction (slope) is shown in Fig. S8A for each tool. For instance, Kallisto counts are related to Reindeer counts by a factor of 108 in average. We found this factor to be related to two phenomena:

- k-mer to fragment or read conversion: given 70 31-mers in each 100-nucleotides read in the CCLE dataset, k-mer counts are expected to be about 140 times higher than fragment counts (each fragment = 2 reads × 70 k-mers),
- query masking: when k-mers in a query are masked, they do not contribute to Reindeer sum counts, whereas other counting methods have no such constraint.

The second point is illustrated by the presence of parallel lines in Fig S9A, suggesting different correction factors for different genes. This is confirmed in Fig S9B, showing slopes of Reindeer vs. Kallisto correlations for independent genes. Most genes have highly correlated counts (*R*^2^ near 1) but each with a distinct slope.

In summary, the correction factor for converting Reindeer sum-counts to read counts from Kallisto or other tools depends both on the library (read size and paired-end status) and the ratio of masked k-mers in each query. A future Reindeer query environment could take these parameters into account and automatically adjust raw k-mer counts in a dataset-and query-independent way. This is not straightforward as it requires transmission of metadata from the indexing and masking steps to the query processing step.

Note that we did not intend to consider here another, non linear, factor of variation, namely that Reindeer counts were obtained based on principal transcript while Kallisto counts were obtained using all isoforms.

### Implementation details

To enable real-time queries with no index loading overhead, we wrote the *rdeer* − *service* tool that pre-loads one or several indexes in memory and runs queries (https://github.com/Bio2M/rdeer-service) on those indexes. *rdeer* −*service* uses a metadata file, *fos.txt*, stored in the same directory as the index, containing names and total numbers of k-mers for each sample. This allows displaying sample names in outputs and computing normalized counts, in the form of counts per billion k-mers.

A *fos.txt* file is not produced by Reindeer index but can be created by the *fos builder* Python script (https://github.com/Transipedia/fos builder). For normalization, *fos builder* uses information from *multiqc*, thus requiring prior run of *fastqc* and *multiqc* on all samples.

## Supplementary Figures

**Figure S1:**
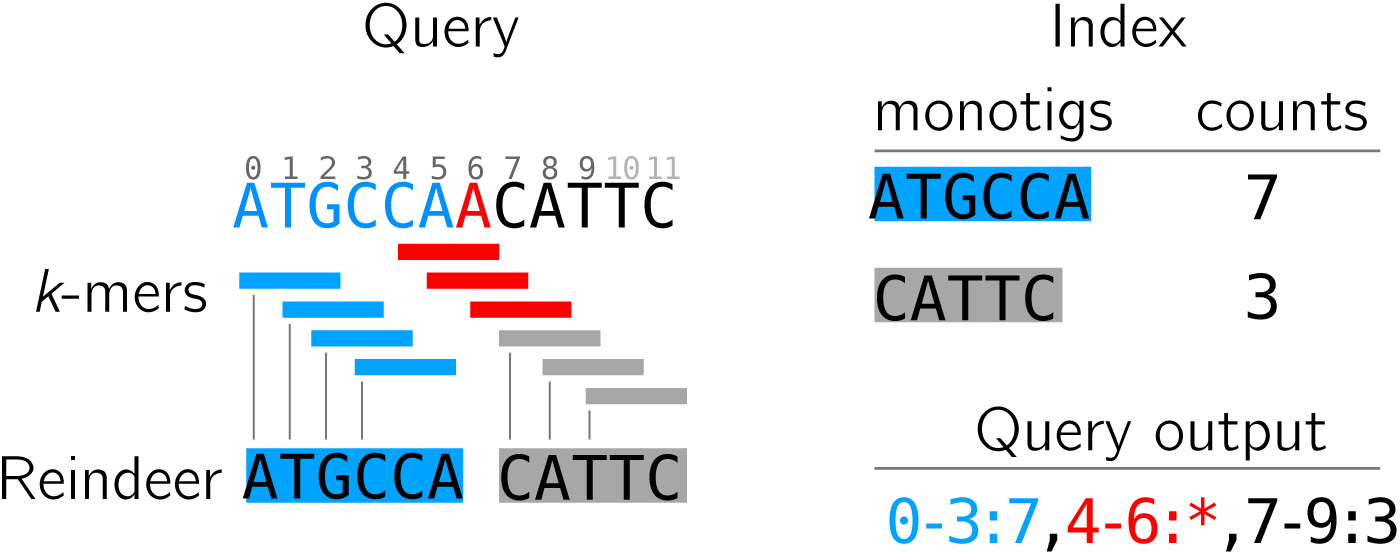
Principle of Reindeer counting. Reindeer natively reports counts for one sample, based on k-mers in the query matched to k-mers in a given monotig index. Monotigs are Reindeer’s subunits that aggregate consecutive k-mers with same counts, they allow to associate a single count to a serie of k-mers. In the example, k-mer size is 3 and the index has two monotigs.

**Figure S2:**
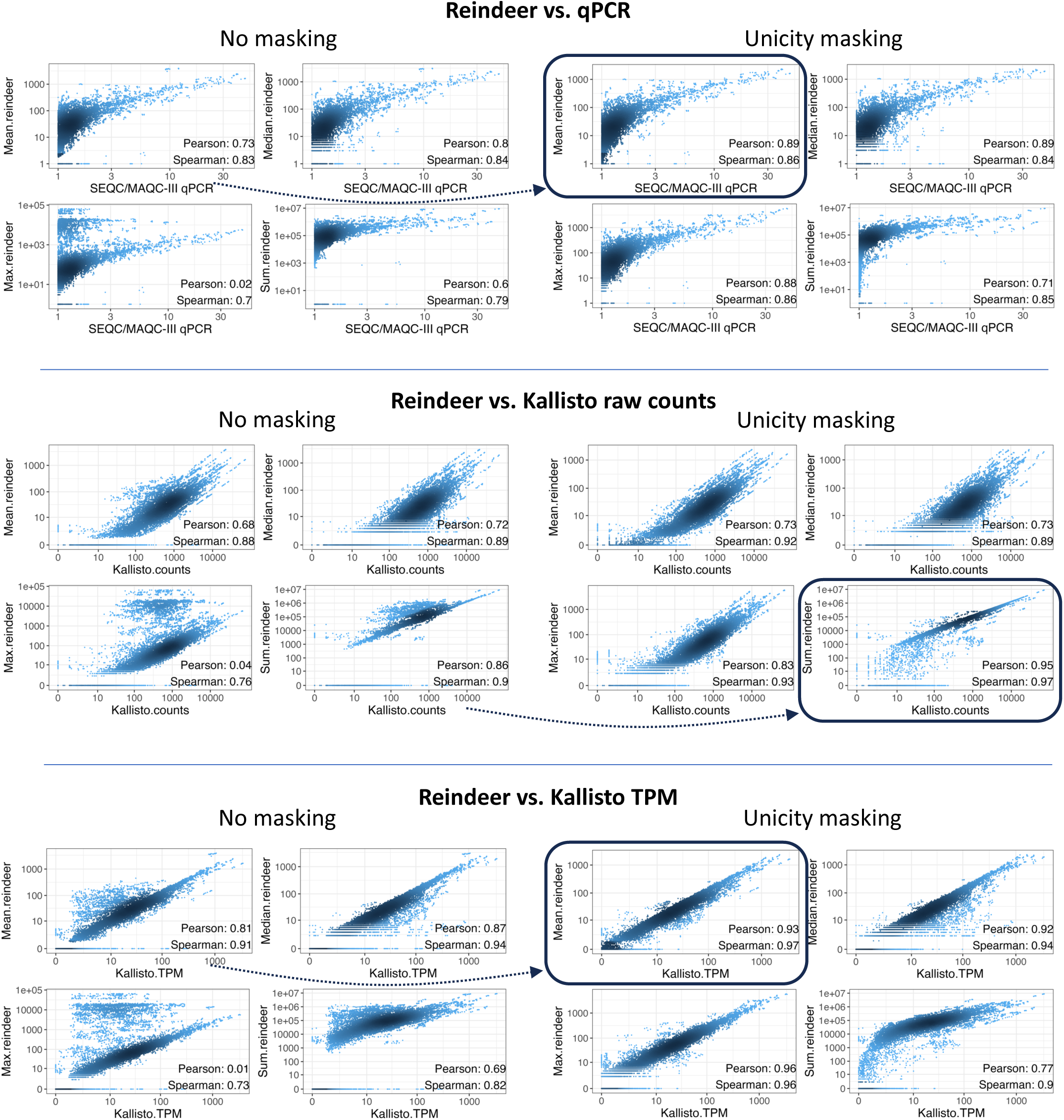
Correlation between Reindeer counts and established count methods, for 1000 genes quantified in 16 reference SEQC/MAQC-III samples. For each comparison, Reindeer counts are computed using one of 4 methods: mean, max, median or sum of monotig counts. On the right side (“masking”) query sequences were first masked through removal of all non-unique k-mers.

**Figure S3:**
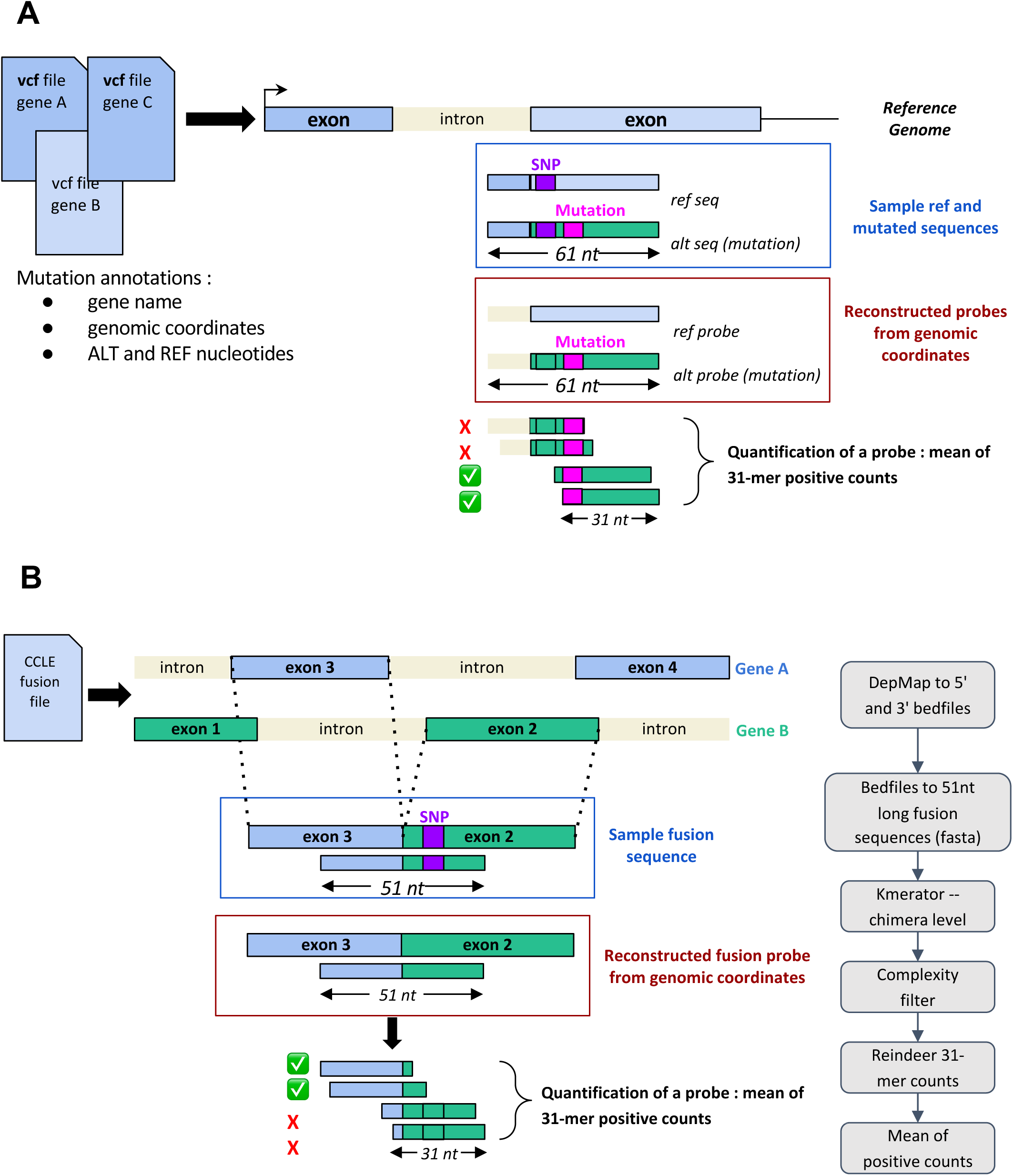
design of mutation/fusion probes reconstructed from genomic coordinates and quantification. **A:** design of 61nt probes for mutation queries. These probes are decomposed into 31-mers for quantification. **B:** design of 51nt probes for fusion queries. These probes are decomposed into 31-mers for quantification. The workflow on the right includes filtering for unicity (Kmerator with --chimera level) and complexity.

**Figure S4:**
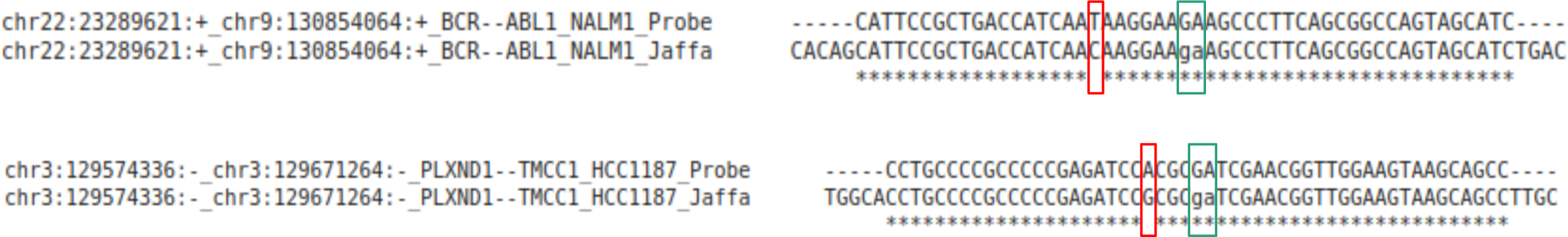
Alignment of the 2 false negative Cosmic fusion probes with one read mapping this fusion described in the Ligea portal (http://hpc-bioinformatics.cineca.it) with the Jaffa algorithm. Red box: variant close to the fusion junction; green box: fusion junction.

**Figure S5:**
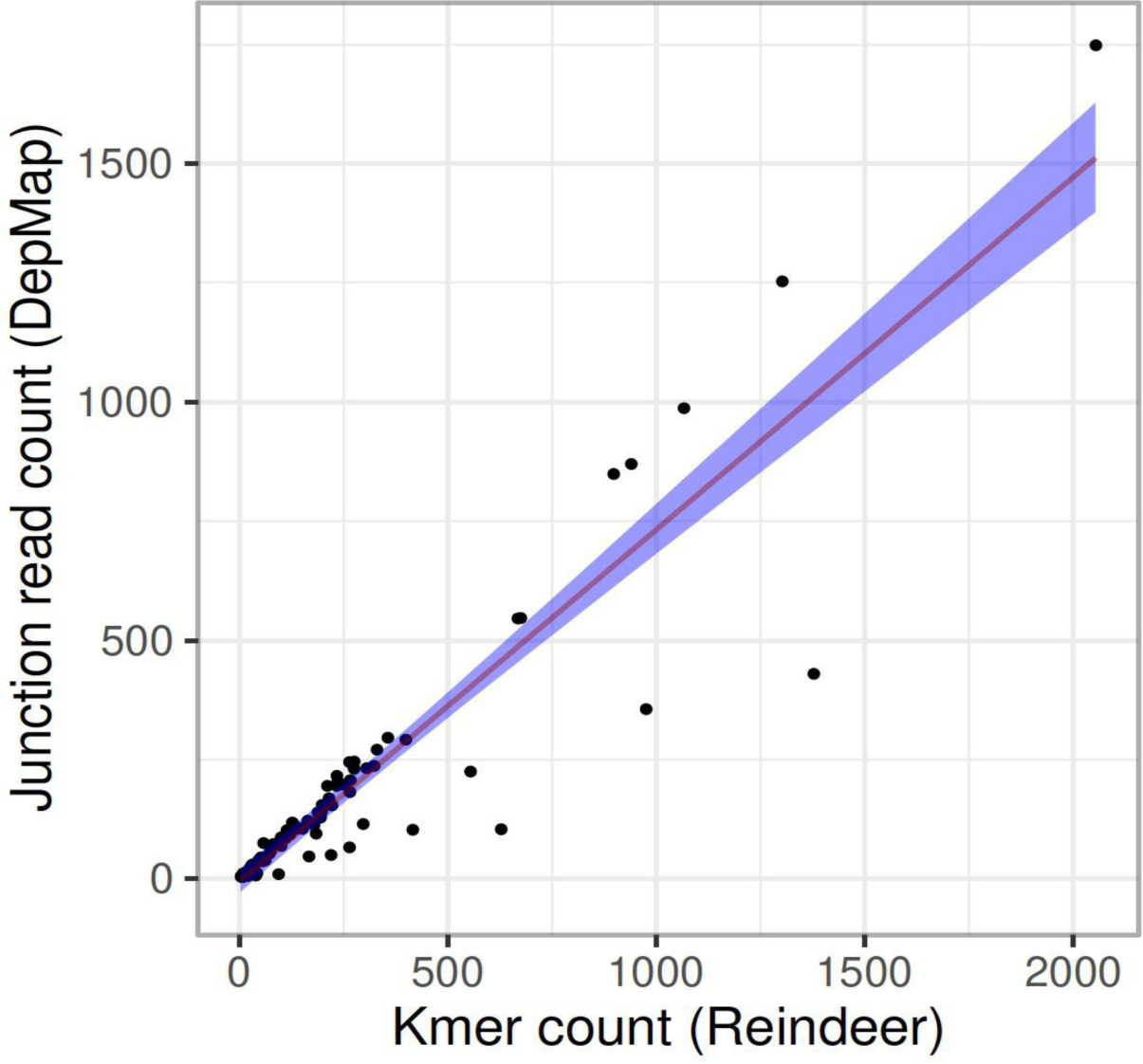
Correlation between Junction read count given by DepMap and Reindeer raw counts for Cosmic fusions.

**Figure S6.**
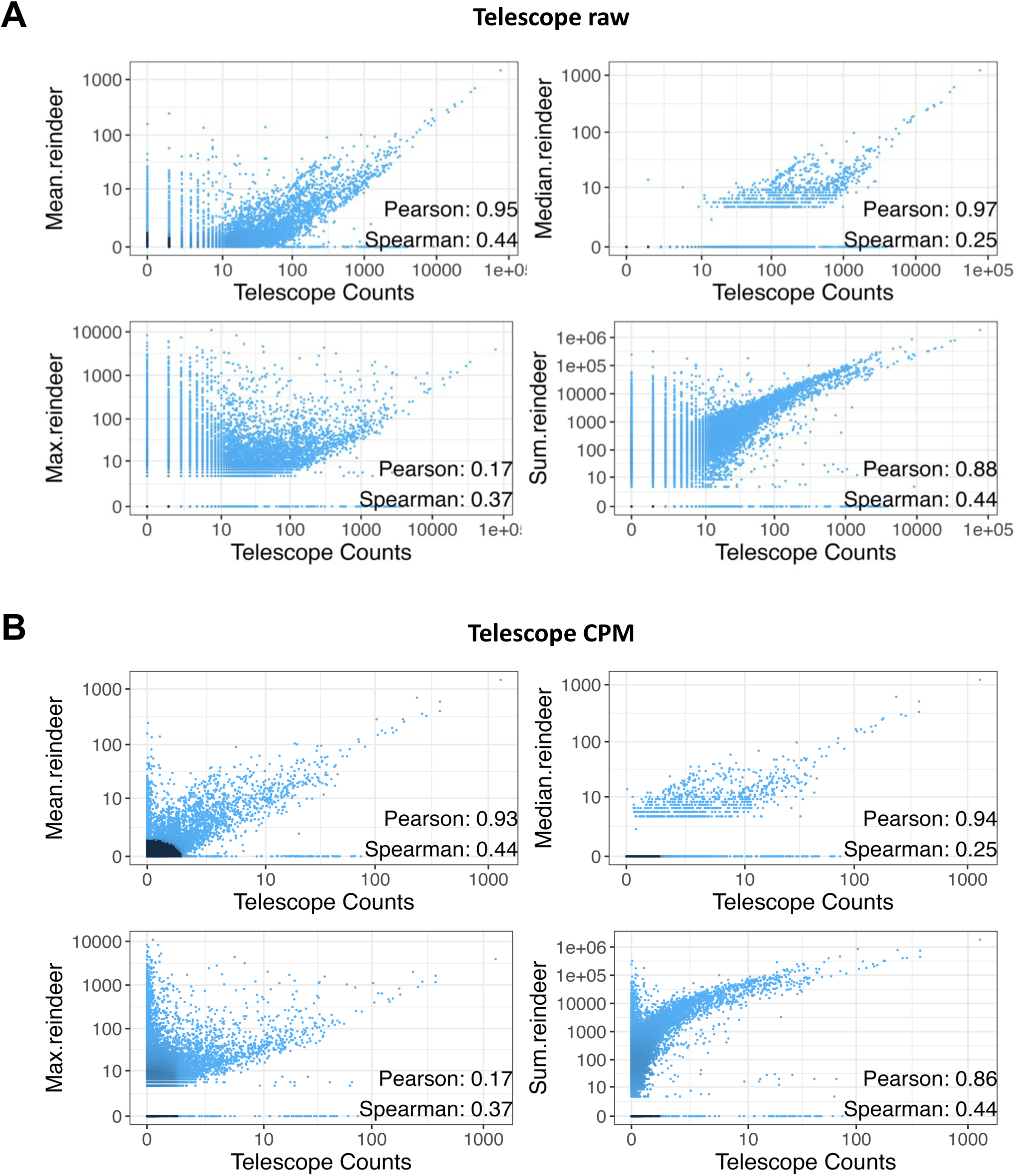
Correlation between Reindeer and Telescope counts. Counts were estimated for 1000 HERV repeats on 56 CCLE colon samples. A: Telescope raw counts. B: Telescope CPM.

**Figure S7:**
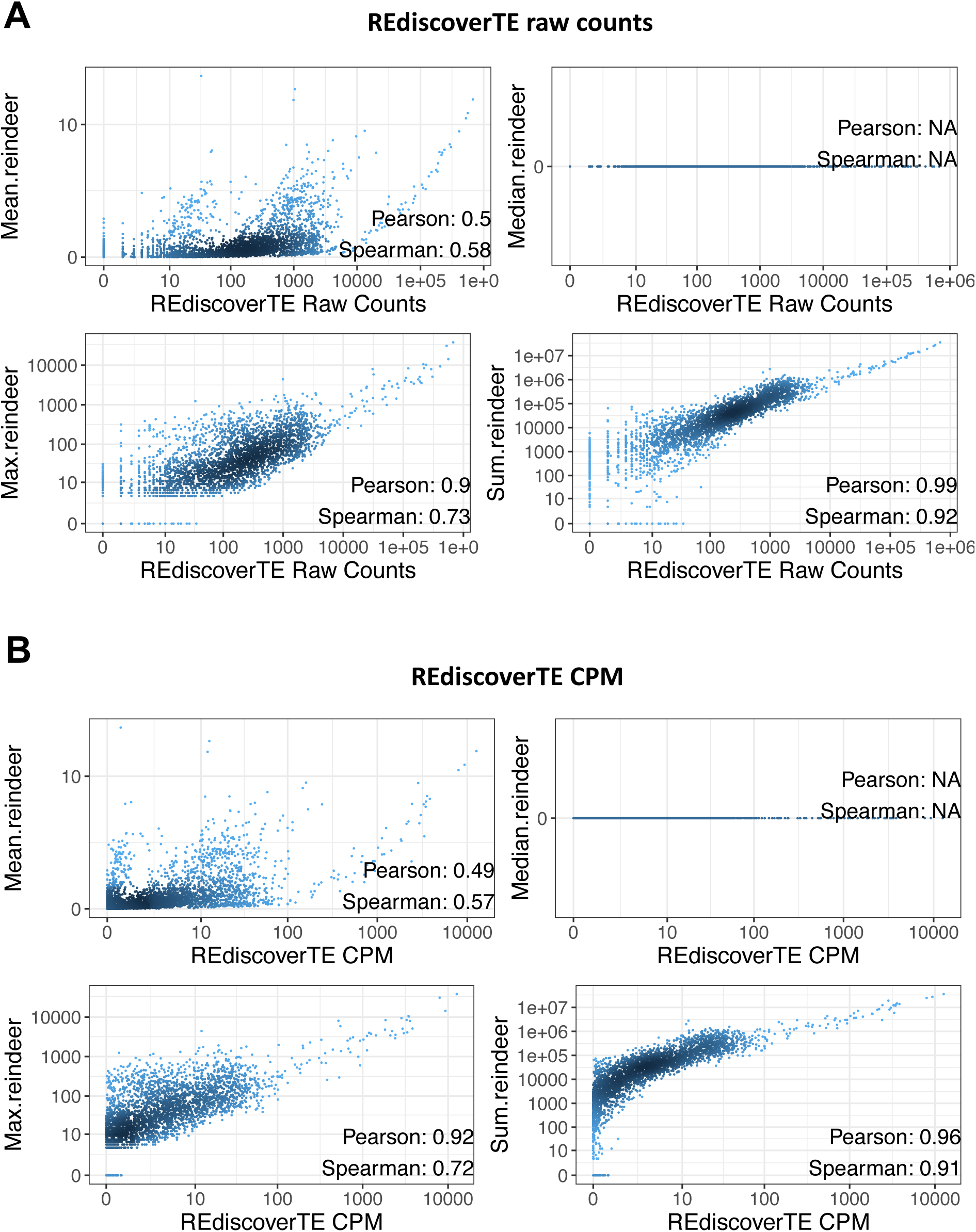
Correlation between Reindeer and REdiscoverTE counts. Counts were estimated for 58 ERV families quantified in 56 CCLE colon samples. **A:** Reindeer vs. REdiscoverTE raw counts. **B:** Reindeer vs. REdiscoverTE CPM. Note that Reindeer median counts for repeat families are always zero. Indeed, each repeat family is composed of hundreds of loci, of which most are silent (count=0), hence the zero median.

**Fig. S8:**
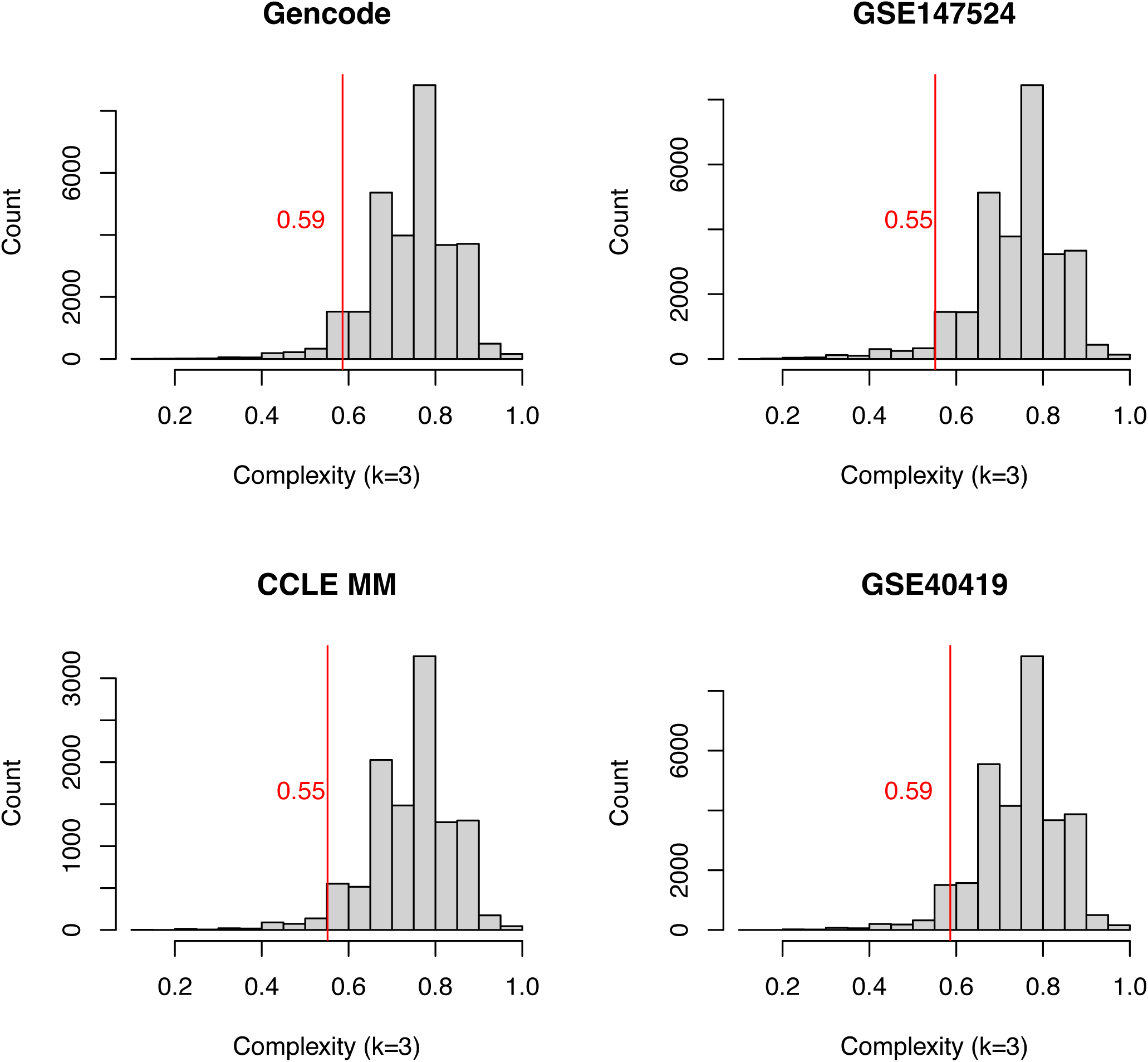
Distribution of 31-mer complexity in four RNA sequence datasets. Complexity is measured for each 31-mer in the dataset as: (number of distinct 3-mers) / (total number of 3-mers). Distribution is shown for Gencode V44, GSE147524 (SRA RNA-seq), GSE40419 (SRA RNA-seq) and CCLE multiple myeloma RNA-seq samples. Fastq datasets were quality-trimmed (Q20) prior to 31-mer extraction. 11,000 to 31,000 31-mers (1/10000^th^) were sampled from each 31-mer table. A vertical red bar is shown at the 5th percentile of each distribution. Based on these observations, the low-complexity threshold for 31-mers was set at 0.55.

**Figure S9:**
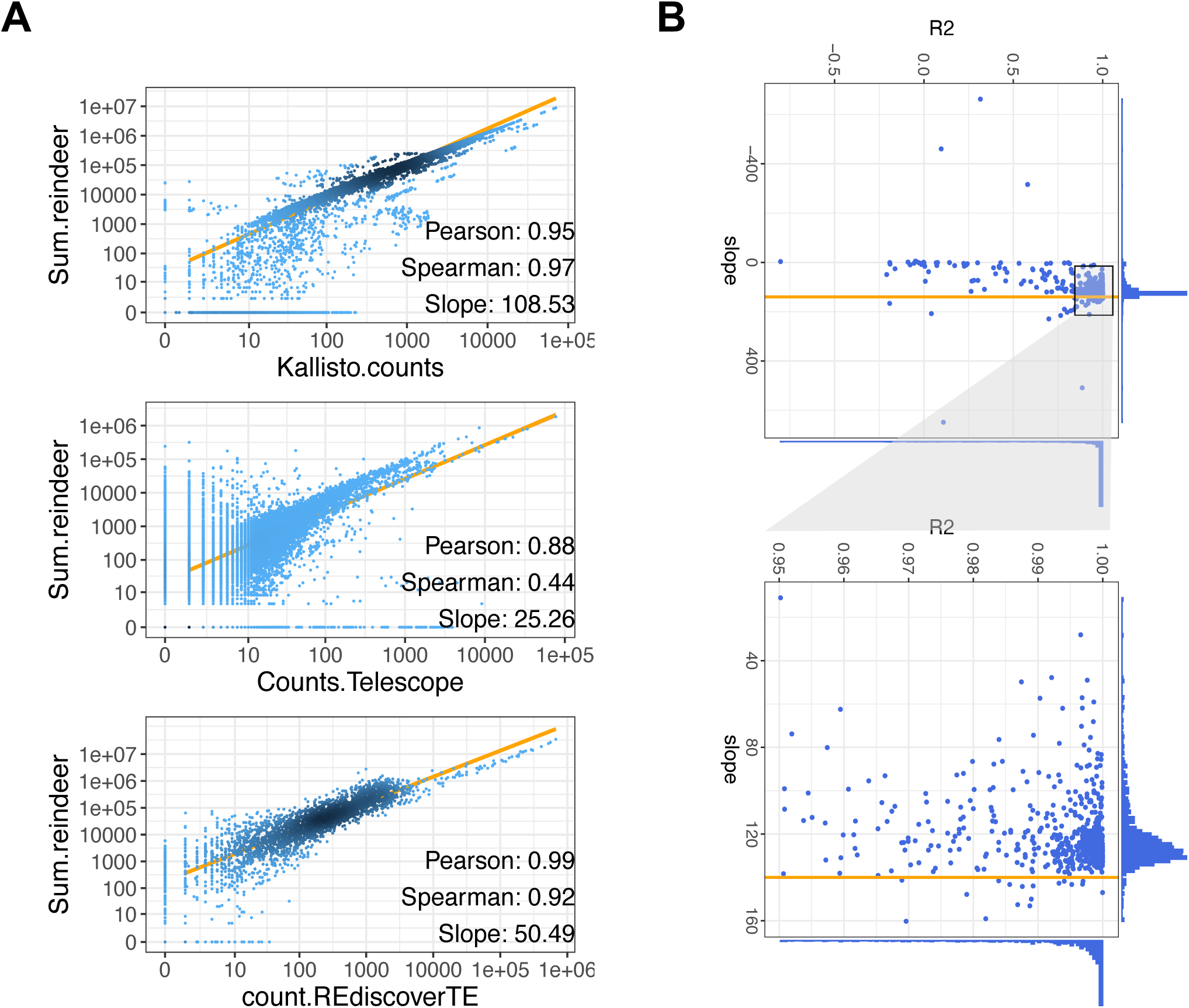
Analysis of correlations between Reindeer sum counts and raw counts from other tools. **A.** Average fitted lines and slopes for Kallisto, Telescope and REdiscoverTE counts (same data as in Fig S1, S6, S7). **B.** Gene by gene slopes and correlation R_2_ for Reindeer vs Kallisto quantificaiton in the MAPQC/SEQC dataset. Each point corresponds to the expression of a single gene measured in 16 conditions. The horizontal line shows a slope of 140. The bottom panel is a zoom on the outlined region on top.

## Notes

https://transipedia.fr

https://github.com/Transipedia/publication-ccle

### Competing Interest Statement

The authors have declared no competing interest.

## References

[1] Lachmann, A. et al. Massive mining of publicly available rna-seq data from human and mouse. Nature communications 9, 1366 (2018).

[2] Clough, E. & Barrett, T. The gene expression omnibus database. Statistical Genomics: Methods and Protocols 93–110 (2016).

[3] Morillon, A. & Gautheret, D. Bridging the gap between reference and real transcriptomes. Genome biology 20, 1–7 (2019).

[4] Wilks, C. et al. recount3: summaries and queries for large-scale rna-seq expression and splicing. Genome biology 22, 1–40 (2021).

[5] Marchet, C. et al. Data structures based on k-mers for querying large collections of sequencing data sets. Genome Research 31, 1–12 (2021).

[6] Darvish, M., Seiler, E., Mehringer, S., Rahn, R. & Reinert, K. Needle: a fast and space-efficient prefilter for estimating the quantification of very large collections of expression experiments. Bioinformatics 38, 4100–4108 (2022).

[7] Karasikov, M., Mustafa, H., Rätsch, G. & Kahles, A. Lossless indexing with counting de bruijn graphs. Genome Research 32, 1754–1764 (2022).

[8] Marchet, C., Iqbal, Z., Gautheret, D., Salson, M. & Chikhi, R. Reindeer: efficient indexing of k-mer presence and abundance in sequencing datasets. Bioinformatics 36, i177–i185 (2020).

9. A comprehensive assessment of rna-seq accuracy, reproducibility and information content by the sequencing quality control consortium. Nature biotechnology 32, 903–914 (2014).

[10] Bray, N. L., Pimentel, H., Melsted, P. & Pachter, L. Near-optimal probabilistic rna-seq quantification. Nature biotechnology 34, 525–527 (2016).

[11] Consortium, C. C. L. E. et al. Genomics of drug sensitivity in cancer consortium. Pharmacogenomic agreement between two cancer cell line data sets. Nature 528, 84–7 (2015).

[12] Tate, J. G. et al. Cosmic: the catalogue of somatic mutations in cancer. Nucleic acids research 47, D941–D947 (2019).

[13] Philippe, N., Salson, M., Commes, T. & Rivals, E. Crac: an integrated approach to the analysis of rna-seq reads. Genome biology 14, 1–16 (2013).

[14] Gillani, R. et al. Gene fusions create partner and collateral dependencies essential to cancer cell survival. Cancer research 81, 3971–3984 (2021).

[15] Davidson, N. M. et al. Jaffal: detecting fusion genes with long-read transcriptome sequencing. Genome biology 23, 1–20 (2022).

[16] Gioiosa, S. et al. Massive ngs data analysis reveals hundreds of potential novel gene fusions in human cell lines. GigaScience 7, giy062 (2018).

[17] Bendall, M. L. et al. Telescope: Characterization of the retrotran-scriptome by accurate estimation of transposable element expression. PLoS computational biology 15, e1006453 (2019).

[18] Kong, Y. et al. Transposable element expression in tumors is associated with immune infiltration and increased antigenicity. Nature communications 10, 5228 (2019).

[19] Patro, R., Duggal, G., Love, M. I., Irizarry, R. A. & Kingsford, C. Salmon provides fast and bias-aware quantification of transcript expression. Nature methods 14, 417–419 (2017).

[20] Alsafadi, S. et al. Cancer-associated sf3b1 mutations affect alternative splicing by promoting alternative branchpoint usage. Nature communications 7, 10615 (2016).

[21] Zhou, Z. et al. The biological function and clinical significance of sf3b1 mutations in cancer. Biomarker research 8, 1–14 (2020).

[22] Alsafadi, S. et al. Genetic alterations of sugp1 mimic mutant-sf3b1 splice pattern in lung adenocarcinoma and other cancers. Oncogene 40, 85–96 (2021).

[23] Riquier, S., et al. Kmerator suite: design of specific k-mer signatures and automatic metadata discovery in large rna-seq datasets. NAR Genomics and Bioinformatics 3, lqab058 (2021).

[24] Chisanga, D., Liao, Y. & Shi, W. Impact of gene annotation choice on the quantification of rna-seq data. BMC bioinformatics 23, 1–21 (2022).

[25] Soneson, C., Love, M. I. & Robinson, M. D. Differential analyses for rna-seq: transcript-level estimates improve gene-level inferences. F1000Research 4 (2015).

[26] Ghandi, M. et al. Next-generation characterization of the cancer cell line encyclopedia. Nature 569, 503–508 (2019).

[27] Kandoth, C. et al. Mutational landscape and significance across 12 major cancer types. Nature 502, 333–339 (2013).

[28] Döhner, H., et al. Diagnosis and management of aml in adults: 2017 eln recommendations from an international expert panel. Blood, The Journal of the American Society of Hematology 129, 424–447 (2017).

[29] Quinlan, A. R. & Hall, I. M. Bedtools: a flexible suite of utilities for comparing genomic features. Bioinformatics 26, 841–842 (2010).

